# Genomics evolutionary history and diagnostics of the *Alternaria alternata* species group including apple and Asian pear pathotypes

**DOI:** 10.1101/534685

**Authors:** Andrew D. Armitage, Helen M. Cockerton, Surapareddy Sreenivasaprasad, James Woodhall, Charles R. Lane, Richard J. Harrison, John P. Clarkson

## Abstract

The *Alternaria* section *alternaria* (*A. alternata* species group) represents a diverse group of saprotroph, human allergens and plant pathogens. *Alternaria* taxonomy has benefited from recent phylogenetic revision but the basis of differentiation between major phylogenetic clades within the group is not yet understood. Furthermore, genomic resources have been limited for the study of host-specific pathotypes. We report near complete genomes of the apple and Asian pear pathotypes as well as draft assemblies for a further 10 isolates representing *Alternaria tenuissima* and *Alternaria arborescens* lineages. These assemblies provide the first insights into differentiation of these taxa as well as allowing the description of effector and non-effector profiles of apple and pear conditionally dispensable chromosomes (CDCs). We define the phylogenetic relationship between the isolates sequenced in this study and a further 23 *Alternaria* spp. based on available genomes. We determine which of these genomes represent MAT1-1-1 or MAT1-2-1 idiomorphs and designate host-specific pathotypes. We show for the first time that the apple pathotype is polyphyletic, present in both the *A. arborescens* and *A. tenuissima* lineages. Furthermore, we profile a wider set of 89 isolates for both mating type idiomorphs and toxin gene markers. Mating-type distribution indicated that gene flow has occurred since the formation of *A. tenuissima* and *A. arborescens* lineages. We also developed primers designed to *AMT14,* a gene from the apple pathotype toxin gene cluster with homologs in all tested pathotypes. These primers allow identification and differentiation of apple, pear and strawberry pathotypes, providing new tools for pathogen diagnostics.

## 1 Introduction

Species within the genus *Alternaria* encompass a range of lifestyles, acting as saprotroph, opportunistic pathogens and host-adapted plant pathogens (Thomma 2003). Large spored species include *A. solani*, a major pathogen of potato, whereas small spored taxa include the *Alternaria alternata* species group (*Alternaria* sect. *alternaria*), which are found ubiquitously in the environment acting as saprotroph and opportunistic necrotrophs. This species group is responsible for opportunistic human infections and a range of host adapted plant diseases.

Taxonomy within this presumed asexual genus has been subject to recent revision (Lawrence et al. 2013). Large spored species can be clearly resolved by standard phylogenetic markers such as ITS and are supported by morphological characters. However, small spored species within the *A. alternata* species group overlap in morphological characters, possess the same ITS haplotype (Kusaba and Tsuge, 1995a) and show low variation in other commonly used barcoding markers (Woudenberg et al. 2015; Lawrence et al. 2013; Armitage et al. 2015). Highly variable phylogenetic markers have provided resolution between groups of isolates that possess morphological patterns typical of descriptions for *A. gaisen*, *A. tenuissima* and *A. arborescens* (Armitage et al. 2015). However, taxonomy of the species group is further complicated by designation of isolates as pathotypes, each able to produce polyketide host-selective toxins (HST) adapted to apple, Asian pear, tangerine, citrus, rough lemon or tomato (Tsuge et al. 2013). Genes involved in the production of these HSTs are located on conditionally dispensable chromosomes (CDCs) (Hatta et al. 2002). CDCs have been estimated to be 1.05 Mb in the strawberry pathotype (Hatta et al. 2002), 1.1 - 1.7 Mb in the apple pathotype (Johnson et al. 2001), 1.1 - 1.9 Mb in the tangerine pathotype (Masunaka et al. 2000; Masunaka et al. 2005) and 4.1 Mb in the pear pathotype (Tanaka et al. 1999; Tanaka and Tsuge 2000). These CDCs are understood to have been acquired through horizontal gene transfer and as such, the evolutionary history of CDCs may be distinct from the core genome.

The polyketide synthase genes responsible for the production of the six HSTs are present in clusters. Some genes within these clusters are conserved between pathotypes (Miyamoto et al. 2009; Hatta et al. 2002), while genes are also present within these clusters that are unique to particular pathotypes (Ajiro et al. 2010; Miyamoto et al. 2010). This is reflected in structural similarities between the pear (AKT) and strawberry (AFT) and tangerine (ACTT) toxins with each containing a 9,10-epoxy-8-hydroxy-9-methyl-decatrienoic acid moiety. In contrast, the toxin produced by the apple pathotype (AMT) does not contain this moiety and is primarily cyclic in structure (Tsuge et al. 2013).

Studies making use of bacterial artificial chromosomes (BAC) have led to the sequencing of toxin gene cluster regions from three apple pathotype isolates (Genbank accessions: AB525198, AB525199, AB525200; unpublished). These sequences are 100-130 Kb in size and contain 17 genes that are considered to be involved in synthesis of the AMT apple toxin (Harimoto et al. 2007). *AMT1*, *AMT2*, *AMT3* and *AMT4* have been demonstrated to be involved in AMT synthesis, as gene disruption experiments have led to loss of toxin production and pathogenicity (Johnson et al. 2000; Harimoto et al. 2007; Harimoto et al. 2008). However, experimental evidence has not been provided to show that the remaining 13 *AMT* genes have a role in toxin production. Four genes present in the CDC for the pear pathotype have been identified and have been named *AKT1*, *AKT2*, *AKT3*, *AKTR-1* (Tanaka et al. 1999; Tanaka & Tsuge 2000; Tsuge et al. 2013) and a further two genes (*AKT4*, *AKTS1*) have been reported (Tsuge et al. 2013).

The toxicity of a HST is not restricted to the designated host for that pathotype. All or some of the derivatives of a toxin may induce necrosis on “non-target” host leaves. For example, AMT from the apple pathotype can induce necrosis on the leaves of Asian pear (Kohmoto et al. 1976). Therefore non-host resistance may be triggered by recognition of non-HST avirulence genes.

*Alternaria spp.* are of phytosanitary importance, with apple and pear pathotypes subject to quarantine regulations in Europe under Annex IIAI of Directive 2000/29/EC as *Alternaria alternata* (non-European pathogenic isolates). As such, rapid and accurate diagnostics are required for identification. Where genes on essential chromosomes can be identified that phylogenetically resolve taxa, then these can be used for identification of quarantine pathogens (Bonants et al. 2010; Quaedvlieg et al. 2012). Regulation and management strategies also need to consider the potential for genetic exchange between species. The *Alternaria* sect. *alternaria* are presumed asexual but evidence has been presented for either the presence of sexuality or a recent sexual past. Sexuality or parasexuality provides a mechanism for reshuffling the core genome associated with CDCs of a pathotype. It is currently unknown whether pathotype identification can be based on sequencing of phylogenetic loci, or whether the use of CDC-specific primers is more appropriate. This is of particular importance for the apple and Asian pear pathotypes due the phytosanitary risk posed by their potential establishment and spread in Europe.

## 2 Methods

Twelve genomes were sequenced, selected from a collection of isolates whose phylogenetic identity was determined in previous work (Armitage et al. 2015). These twelve isolates were three *A. arborescens* clade isolates (*FERA 675*, *RGR 97.0013* and *RGR 97.0016*), four *A. tenuissima* clade isolates (*FERA 648*, *FERA 1082*, *FERA 1164* and *FERA 24350*), three *A. tenuissima* clade apple pathotype isolates (*FERA 635*, *FERA 743*, *FERA 1166* and *FERA 1177*) and one *A. gaisen* clade pear pathotype isolate (*FERA 650*).

### 2.1 DNA and RNA extraction and sequencing

Apple pathotype isolate *FERA 1166* and Asian pear pathotype isolate *FERA 650* were sequenced using both Illumina and nanopore MinION sequencing technologies and the remaining ten isolates were sequenced using Illumina sequencing technology. For both illumina and MinION sequencing, DNA extraction was performed on freeze dried mycelium grown in PDB for 14 days.

High molecular weight DNA was extracted for MinION sequencing using the protocol of Schwessinger & McDonald (2017), scaled down to a starting volume of 2 ml. This was followed by phenol-chloroform purification and size selection to a minimum of 30 Kb using a Blue Pippin. The resulting product was concentrated using ampure beads before library preparation was performed using a Rapid Barcoding Sequencing Kit (SQK-RBK001) modified through exclusion of LLB beads. Sequencing was performed on a Oxford Nanopore GridION generating an additional 39 and 44 times coverage of sequence data for isolates *FERA 1166* and *FERA 650* respectively.

gDNA for illumina sequencing of isolate *FERA 1166* was extracted using a modified CTAB protocol (Li *et al.*, 1994). gDNA for illumina sequencing of the eleven other isolates was extracted using a Genelute Plant DNA Miniprep Kit (Sigma) using the manufacturers protocol with the following modifications: the volume of lysis solutions (PartA and PartB) were doubled; an RNase digestion step was performed as suggested in the manufacturer's protocol; twice the volume of precipitation solution was added; elution was performed using elution buffer EB (Qiagen). A 200 bp genomic library was prepared for isolate *FERA 1166* using a TrueSeq protocol (TrueSeq Kit, Illumina) and sequenced using 76 bp paired-end reads on an Illumina GA2 Genome Analyser. Genomic libraries were prepared for the other eleven isolates using a Nextera Sample Preparation Kit (Illumina) and libraries sequenced using a MiSeq Benchtop Analyser (Illumina) using 250 bp, paired end reads.

RNAseq was performed to aid training of gene models. mRNA was extracted from isolates *FERA 1166* and *FERA 650* grown in full strength PDB, 1% PDB, Potato Carrot Broth (PCB) and V8 juice broth (V8B). The protocol for making PCB and V8B was as described in (Simmons, 2007)for making Potato Carrot Agar and V8 juice agar, with the exception that agar was not added to the recipe. Cultures were grown in conical flasks containing 250 mls of each liquid medium for 14 days. mRNA extraction was performed on freeze dried mycelium using the RNeasy Plant RNA extraction Kit (Qiagen). Concentration and quality of mRNA samples were assessed using a Bioanalyzer (Agilent Technologies). mRNA from the sample grown in 1% PDB for isolate *FERA 650* showed evidence of degradation and was not used further. Samples were pooled from growth mediums for each isolate and 200 bp cDNA libraries prepared using a TrueSeq Kit (Illumina). These libraries were sequenced in multiplex on a MiSeq (Illumina) using 200 bp paired end reads.

### 2.2 Genome assembly and annotation

*De-novo* genome assembly was performed for all 12 isolates. Assembly for isolate *FERA 650* was generated using SMARTdenovo (https://github.com/ruanjue/smartdenovo, February 26, 2017 github commit), whereas assembly for isolate *FERA 1166* was generated by merging a SMARTdenovo assembly with a MinION-Illumina hybrid SPAdes v3.9.0 assembly using quickmerge v0,2 (Antipov et al. 2016, Chakraborty et al. 2016). Prior to assembly, adapters were removed from MinION reads using Porechop v0.1.0 and reads were further trimmed and corrected using Canu v1.6 (Ruan 2016; Koren et al. 2017). Following initial assembly, contigs were corrected using MinION reads through ten rounds of Racon (May 29, 2017 github commit) correction (Vaser et al. 2017)and one round of correction using MinION signal information with nanopolish (v0.9.0) (Loman et al. 2015). Final correction was performed through ten rounds of Pilon v1.17 (Walker et al. 2014) using Illumina sequence data. Assemblies for the ten isolates with Illumina-only data were generated using SPAdes v.3.9.0 (Bankevich et al. 2012). Assembly quality statistics were summarized using Quast v.4.5 (Gurevich et al. 2013). Single copy core Ascomycete genes were identified within the assembly using BUSCO v3 and used to assess assembly completeness (Simão et al. 2015). RepeatModeler, RepeatMasker and TransposonPSI were used to identify repetitive and low complexity regions (http://www.repeatmasker.org, http://transposonpsi.sourceforge.net). Visualisation of whole genome alignments between *FERA 1166* and *FERA 650* was performed using circos v0.6 (Krzywinski et al. 2009), following whole genome alignment using the nucmer tool as part of the MUMmer package v4.0 (Marçais *et al.* 2018).

Gene prediction was performed on softmasked genomes using Braker1 v.2 (Hoff et al. 2016), a pipeline for automated training and gene prediction of AUGUSTUS 3.1 (Stanke & Morgenstern 2005). Additional gene models were called in intergenic regions using CodingQuarry v.2 (Testa et al. 2015). Braker1 was run using the “fungal” flag and CodingQuarry was run using the “pathogen” flag. RNAseq data generated from *FERA 1166* and *FERA 650* were aligned to each genome using STAR v2.5.3a (Dobin et al. 2013), and used in the training of Braker1 and CodingQuarry gene models. Orthology was identified between the 12 predicted proteomes using OrthoMCL v.2.0.9 (Li et al. 2003) with an inflation value of 5. Orthology was identified between the 12 predicted proteomes using OrthoMCL v.2.0.9 (Li et al. 2003) with an inflation value of 5.

Draft functional genome annotations were determined for gene models using InterProScan-5.18-57.0 (Jones et al. 2014) and through identifying homology (BLASTP, E-value > 1×10^−100^) between predicted proteins and those contained in the March 2018 release of the SwissProt database (Bairoch and Apweiler 2000). Putative secreted proteins were identified through prediction of signal peptides using SignalP v.4.1 and removing those predicted to contain transmembrane domains using TMHMM v.2.0 (Käll et al. 2004; Krogh et al. 2001). Additional programs were used to provide evidence of effectors and pathogenicity factors. EffectorP v1.0 was used to screen secreted proteins for characteristics of length, net charge and amino acid content typical of fungal effectors (Sperschneider et al. 2016). Secreted proteins were also screened for carbohydrate active enzymes using HMMER3 (Mistry et al. 2013) and HMM models from the dbCAN database (Huang et al. 2018). DNA binding domains associated with transcription factors (Shelest 2017) were identified along with two additional fungal-specific transcription factors domains (IPR007219 and IPR021858). Annotated assemblies were submitted as Whole Genome Shotgun projects to DDBJ/ENA/Genbank (Table 1).

**Table 1.** NCBI biosample and genome accession numbers of data generated in this study.

| Isolate | BioSample | Accession |
| --- | --- | --- |
| *FERA 675* | SAMN06205217 | PDUP01000000 |
| *RGR 97.0013* | SAMN06205218 | PDWY01000000 |
| *RGR 97.0016* | SAMN06205219 | PEJP01000000 |
| *FERA 650* | SAMN06205220 | PDWZ02000000 |
| *FERA 1082* | SAMN06205221 | PDXA01000000 |
| *FERA 1164* | SAMN06205222 | PDXB01000000 |
| *FERA 1166* | SAMN06205223 | PDXC01000000 |
| *FERA 1177* | SAMN06205224 | PDXD01000000 |
| *FERA 24350* | SAMN06205225 | PDXE01000000 |
| *FERA 635* | SAMN06205226 | PDXF01000000 |
| *FERA 648* | SAMN06205227 | PDXG01000000 |
| *FERA 743* | SAMN06205228 | PDXH01000000 |

### 2.3 Phylogenetics

BUSCO hits of single copy core ascomycete genes to assemblies were extracted and retained if a single hit was found in all of the 12 sequenced genomes and 23 publically available *Alternaria* spp. genomes from the Alternaria genomes database (Dang et al. 2015). Sequences from the resulting hits of 500 loci were aligned using MAFFT v6.864b (Katoh and Standley 2013), before alignments were trimmed using trimAl v1.4.1 (Capella-Gutiérrez et al. 2009), and trees calculated for each locus using RAxML v8.1.17 (Liu et al. 2011). The most parsimonious tree from each RAxML run was used to determine a single consensus phylogeny of the 500 loci using ASTRAL v5.6.1 (Zhang et al. 2018). The resulting tree was visualised using the R package GGtree v1.12.4 (Yu et al. 2016).

### 2.4 CDC identification

Contigs unique to apple and pear pathotypes were identified through read alignment to assembled genomes. Read alignment was performed using Bowtie2 (Langmead and Salzberg 2012), returning a single best alignment for each paired read. Read coverage was quantified from these alignments using Samtools (Li et al. 2009).

### 2.5 Toxin-synthesis genes in *Alternaria* genomes

Sequence data for 40 genes located in *A. alternata* HST gene clusters were downloaded from Genbank. BLASTn searches were performed for all 40 gene sequences against one another to identify homology between these sequences. Genes were considered homologous where they had > 70 % identical sequences over the entire query length, and an e-value of 1×10^−30^. tBLASTx was used to search for the presence of these genes in assemblies.

### 2.6 Signatures of genetic exchange

Mating type idiomorphs present in publicly available genomes were identified using BLASTn searches. A wider assessment within 89 characterised *Alternaria* isolates (Armitage et al. 2015) was undertaken using specific PCR primers (Arie et al. 2000). PCR primers (AAM1-3: 5’-TCCCAAACTCGCAGTGGCAAG-3’; AAM1-3: 5’-GATTACTCTTCTCCGCAGTG-3; M2F: 5’-AAGGCTCCTCGACCGATGAA-3; M2R: 5’-CTGGGAGTATACTTGTAGTC-3) were run in multiplex with PCR reaction mixtures consisting of 10 μl redtaq (REDTaq ReadyMix PCR Reaction Mix, Sigma-Aldrich), 2 μl DNA, 1 μl of each primer (20 μM), and 4 μl purified water (Sigma-Aldrich). PCR reaction conditions comprised of an initial 60 second denaturing step at 94°C followed by 30 cycles of a melting step of 94°C for 30 seconds, an annealing step at 57°C for 30 seconds, and an extension step at 72°C for 60 seconds, these cycles were followed by a final extension step at 72°C for 420 seconds. MAT1-1-1 or MAT1-2-1 idiomorphs were determined through presence of a 271 or 576 bp product following gel electrophoresis, respectively.

### 2.7 PCR screens for apple and pear toxin-synthesis genes

A set of 90 previously characterised isolates was used to further investigate the distribution of pathotypes throughout the *A. alternata* species group. PCR primers were designed for the amplification of three genes (*AMT4*, *AMT14*, *AKT3*) located within CDC gene clusters involved in toxin synthesis. Primers for *AMT4* were designed to amplify apple pathotype isolates, *AKT3* to amplify pear pathotype isolates and *AMT14* to identify both apple and pear pathotype isolates. These primers were then used to screen isolates for the presence of these genes in 30 cycles of PCR using 0.25 ul Dream taq, 1 ul of 10x PCR buffer, 1 ul of dNTPs, 1 ul of gDNA, 1 ul of each primer (5 μM), and 4.75 ul purified water (Sigma-Aldrich). PCR products were visualised using gel electrophoresis and amplicon identity confirmed through Sanger sequencing. Primers AMT4-EMR-F (5’-CTCGACGACGGTTTGGAGAA-3) and AMT4-EMR-R (5’-TTCCTTCGCATCAATGCCCT-3) were used for amplification of AMT4. Primers AKT3-EMR-F (5’-GCAATGGACGCAGACGATTC-3) and AKT3-EMR-R (5’-CTTGGAAGCCAGGCCAACTA-3) were used for amplification of *AKT3*. Primers AMT14-EMR-F (5’-TTTCTGCAACGGCGKCGCTT-3) and AMT14-EMR-R (5’-TGAGGAGTYAGACCRGRCGC-3) were used for amplification of *AMT14*. PCR reaction conditions were the same as described above for mating type loci, but with annealing performed at 66°C for all primer pairs.

### 2.8 Virulence assay

Pathogenicity assays were performed on apple *cv.* Spartan and *cv.* Bramley’s seedling to determine differences in isolate virulence between *A. tenuissima* isolates possessing the apple pathotype CDC (*FERA 635*, *FERA 743* or *FERA 1166*) and non-pathotype isolates lacking the CDC (*FERA 648*, *FERA 1082* or *FERA 1164*). Detailed pathogenicity assay methods are provided in supplementary information 5. Briefly, leaves were inoculated with 10 ul of 1×10^5^ spore suspensions at six points and the number of leaf spots counted at 14 days post inoculation. One isolate was infected per leaf, with 10 replicates per cultivar. Binomial regression using a generalised linear model (GLM) was used to analyse the number of resulting lesions per leaf.

Unfolded adult apple leaves, less than 10 cm in length were cut from young (less than 12 months old) apple *cv.* Spartan trees or *cv.* Bramley’s seedling trees. These were quality-checked to ensure that they were healthy and free from disease. Leaves were grouped by similar size and age and organised into ten experimental replicates of nine leaves. Leaves placed in clear plastic containers, with the abaxial leaf surface facing upwards. The base of these boxes was lined with two sheets of paper towel, and wetted with 50 mls of sterile distilled water (SDW). The cultivars were assessed in two independent experiments.

Spore suspensions were made by growing *A. alternata* isolates on 1% PDA plates for four weeks at 23 °C before flooding the plate with 2 mls of SDW, scraping the plate with a disposable L-shaped spreader. Each leaf was inoculated with 10 ml of 1×10^5^ spores.ml^−1^ *A. alternata* spore suspension or 10 ml of sterile-distilled water at six points on the abaxial leaf surface. Of the nine leaves in each box, three leaves were inoculated with a spore suspensions from isolates carrying apple pathotype CDC, three leaves were inoculated with non-pathotype isolates lacking the CDC, and three leaves were inoculated with SDW. Following inoculation, each container was sealed and placed in plastic bags to prevent moisture loss. Boxes were then kept at 23 °C with a 12 hr. light / 12 hr. dark cycle.

## 3 Results

### 3.1 Generation of near-complete genomes for the apple and pear pathotype using MinION sequencing

Assemblies using nanopore long-read sequence data for the apple pathotype isolate *FERA 1166*, and pear pathotype isolate *FERA 650* were highly contiguous, with the former totaling 35.7 Mb in 22 contigs and the latter totaling 34.3 Mb in 27 contigs (Table 2). Whole genome alignments of these assemblies to the 10 chromosomes of *A. solani* showed an overall macrosynteny between genomes (Fig. 1), but with structural rearrangement of apple pathotype chromosomes in comparison to *A. solani* chromosomes 1 and 10. The Asian pear pathotype had distinct structural rearrangements in comparison to *A. solani*, chromosomes 1 and 2 (Fig. 1). Scaffolded contigs of *FERA 1166* spanned the entire length of *A. solani* chromosomes 2, 3, 6, 8 and 9 and chromosomes 4, 5, 6 and 10 for *FERA 650* (Fig. 1). Interestingly, sites of major structural rearrangements within *A. solani* chromosome 1 were flanked by telomere-like TTAGGG sequences.

**Table 2.** Assembly and gene prediction statistics for genomes from three A. alternata species group lineages, including apple and Asian pear pathotype isolates. Number of genes predicted to encode secreted proteins, secreted effectors (EffectorP) and secreted carbohydrate active enzymes (CAZymes) are shown as well as the total number of secondary metabolite clusters in the genome. The percentage of 1315 conserved ascomycete genes that were identified as complete and present in a single copy within assemblies or gene models are shown.

| Organism | *A. gaisen* | *A. tenuissima* |  |  |  |  |  |  |  | *A. arborescens* |  |  |
| --- | --- | --- | --- | --- | --- | --- | --- | --- | --- | --- | --- | --- |
| Pathotype | pear | apple |  |  |  | non-pathotype |  |  |  | non-pathotype |  |  |
| Isolate | FERA | FERA | FERA | FERA | FERA | FERA | FERA | FERA | FERA | FERA | RGR | RGR |
|  | 650 | 1166 | 635 | 743 | 1177 | 648 | 1082 | 1164 | 24350 | 675 | 97.0013 | 97.0016 |
| Assembly size (Mb) | 34.3 | 35.7 | 36.1 | 35.9 | 35.6 | 33.5 | 33.9 | 34.7 | 33.0 | 33.9 | 33.8 | 33.8 |
| Contigs | 27 | 22 | 912 | 788 | 735 | 124 | 345 | 250 | 167 | 325 | 287 | 339 |
| Largest contig (Mb) | 6.26 | 3.90 | 1.43 | 3.88 | 4.61 | 3.50 | 1.72 | 2.04 | 1.90 | 2.09 | 1.61 | 1.06 |
| N50 (Kb) | 2,110 | 1,584 | 319 | 622 | 537 | 1,341 | 547 | 709 | 608 | 862 | 685 | 528 |
| Repeatmasked (Kb) | 749 | 1,268 | 953 | 869 | 865 | 658 | 628 | 983 | 466 | 853 | 955 | 784 |
| Repeatmasked (%) | 2.18 | 3.55 | 2.64 | 2.42 | 2.43 | 1.96 | 1.85 | 2.83 | 1.41 | 2.51 | 2.83 | 2.32 |
| Conserved Ascomycete genes in genome (%) | 98.7 | 99.0 | 98.8 | 98.9 | 98.8 | 99.0 | 98.6 | 98.8 | 99.0 | 98.6 | 98.6 | 98.7 |
| Total genes | 13169 | 13576 | 13733 | 13707 | 13580 | 12757 | 13028 | 13114 | 12806 | 12896 | 12766 | 12820 |
| Total proteins | 13220 | 13633 | 13812 | 13776 | 13647 | 12798 | 13091 | 13169 | 12856 | 12936 | 12813 | 12863 |
| Conserved Ascomycete genes in gene models (%) | 98.7 | 99.0 | 98.1 | 98.2 | 97.7 | 98.3 | 98.2 | 98.5 | 98.4 | 97.7 | 98.1 | 98.1 |
| Secreted proteins | 1208 | 1251 | 1261 | 1247 | 1270 | 1228 | 1246 | 1235 | 1225 | 1199 | 1189 | 1185 |
| Secreted EffectorP proteins | 246 | 248 | 264 | 263 | 268 | 233 | 252 | 252 | 241 | 236 | 229 | 229 |
| Secreted CAZYme proteins | 383 | 389 | 397 | 390 | 397 | 392 | 401 | 385 | 391 | 375 | 372 | 382 |
| Secondary metabolite clusters | 30 | 34 | 36 | 36 | 30 | 28 | 30 | 29 | 26 | 28 | 28 | 30 |

**Figure 1.**
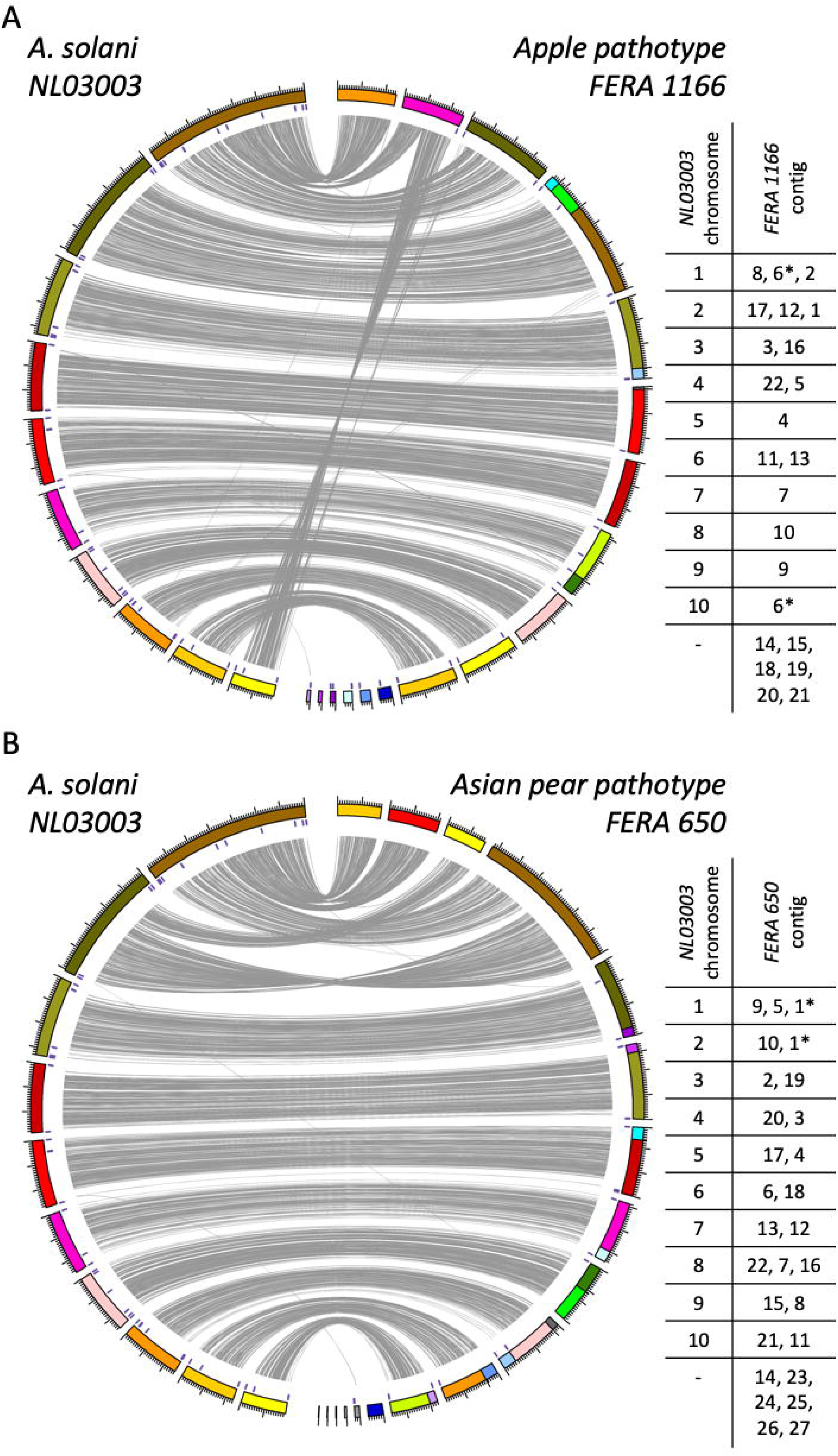
Genome alignment between the reference *A. solani* genome and long-read assemblies of *A. alternata* apple (A) and Asian pear (B) pathotypes. Links are shown between aligned regions. Locations of telomere repeat sequences are marked within assembled contigs. Contig order in reference to *A. solani* chromosomes is summarized, with those contigs displaying evidence of structural rearrangement marked with an asterisk.

Genome assembly of 10 Illumina sequenced isolates yielded assemblies of a similar total size to MinION assemblies (33.9 - 36.1 Mb) but fragmented into 167-912 contigs. Assembled genomes were repeat sparse, with 1.41 - 2.83 % of genomes repeat masked (Table 2). Genome assemblies of *A. arborescens* isolates (33.8 - 33.9 Mb), were of similar total size to non-pathotype *A. tenuissima* isolates and had similar repetitive content (2.51 – 2.83 and 1.41 – 2.83 % respectively). Despite this, identification of transposon families in both genomes showed expansion of DDE (T_5df_ = 5.36, P > 0.01) and gypsy (T_5df_ = 6.35, P > 0.01,) families in *A. arborescens* genomes (Fig. 2).

**Figure 2.**
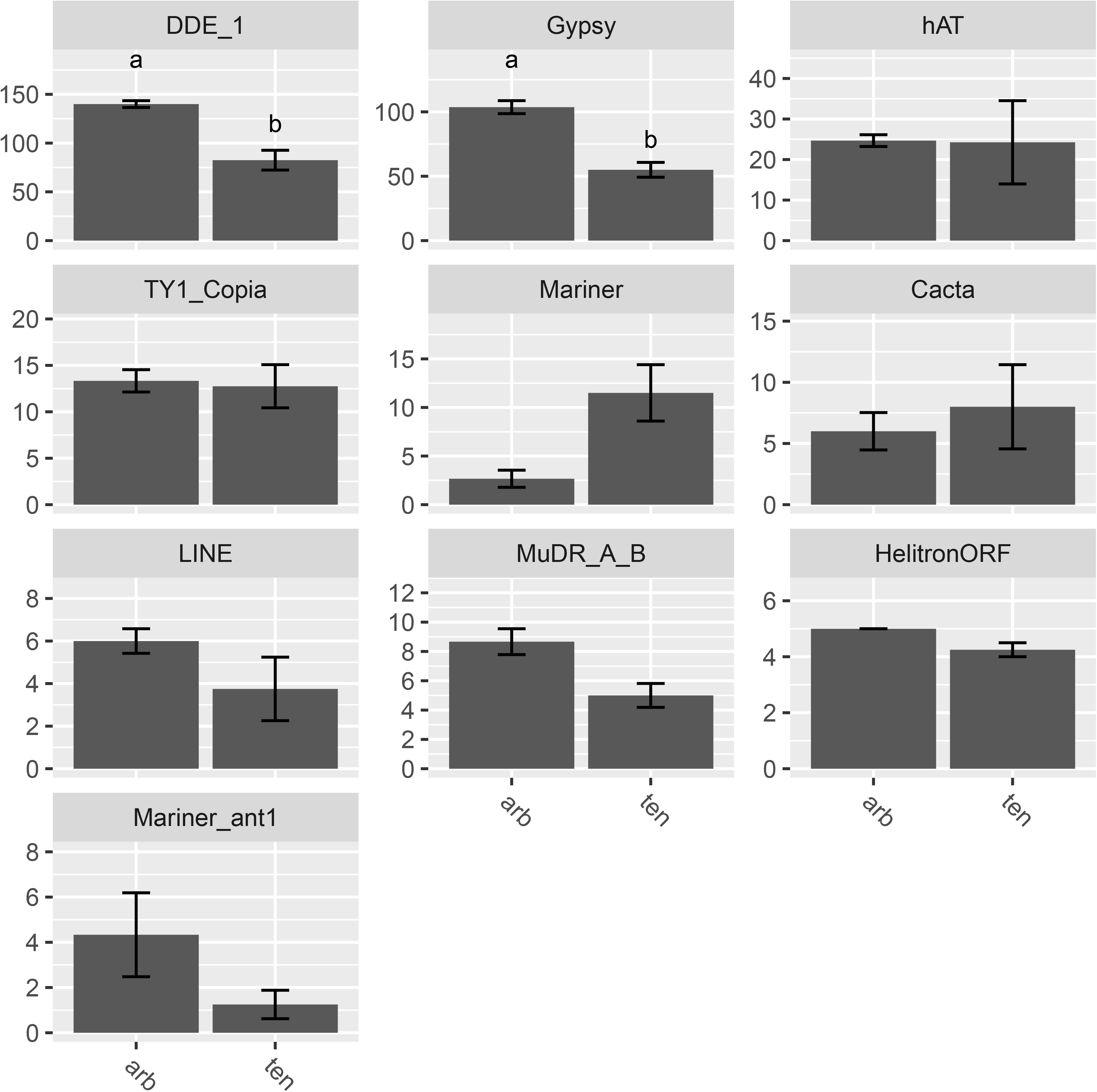
Distinct DDE and gypsy family transposon families between genomes of *A. arborescens* (arb) and *A. tenuissima* (ten) clade isolates. Numbers of identified transposons are also shown for hAT, TY1 copia, mariner, cacta, LINE, MuDR / Mu transposases, helitrons and the Ant1-like mariner elements.

### 3.2 Phylogeny of sequenced isolates

The relationship between the 12 sequenced isolates and 23 *Alternaria* spp. with publically available genomes was investigated through phylogenetic analysis of 500 shared core ascomycete genes. *A. pori* and *A. destruens* genomes were excluded from the analysis due to low numbers of complete single copy ascomycete genes being found in their assemblies (Supp. information 1). The 12 sequenced isolates were distributed throughout *A. gaisen*, *A. tenuissima* and *A. arborescens* clades (Fig. 3). The resulting phylogeny (Fig. 3), formed the basis for later assessment of CDC presence and mating type distribution among newly sequenced and publicly available genomes, as discussed below.

**Figure 3.**
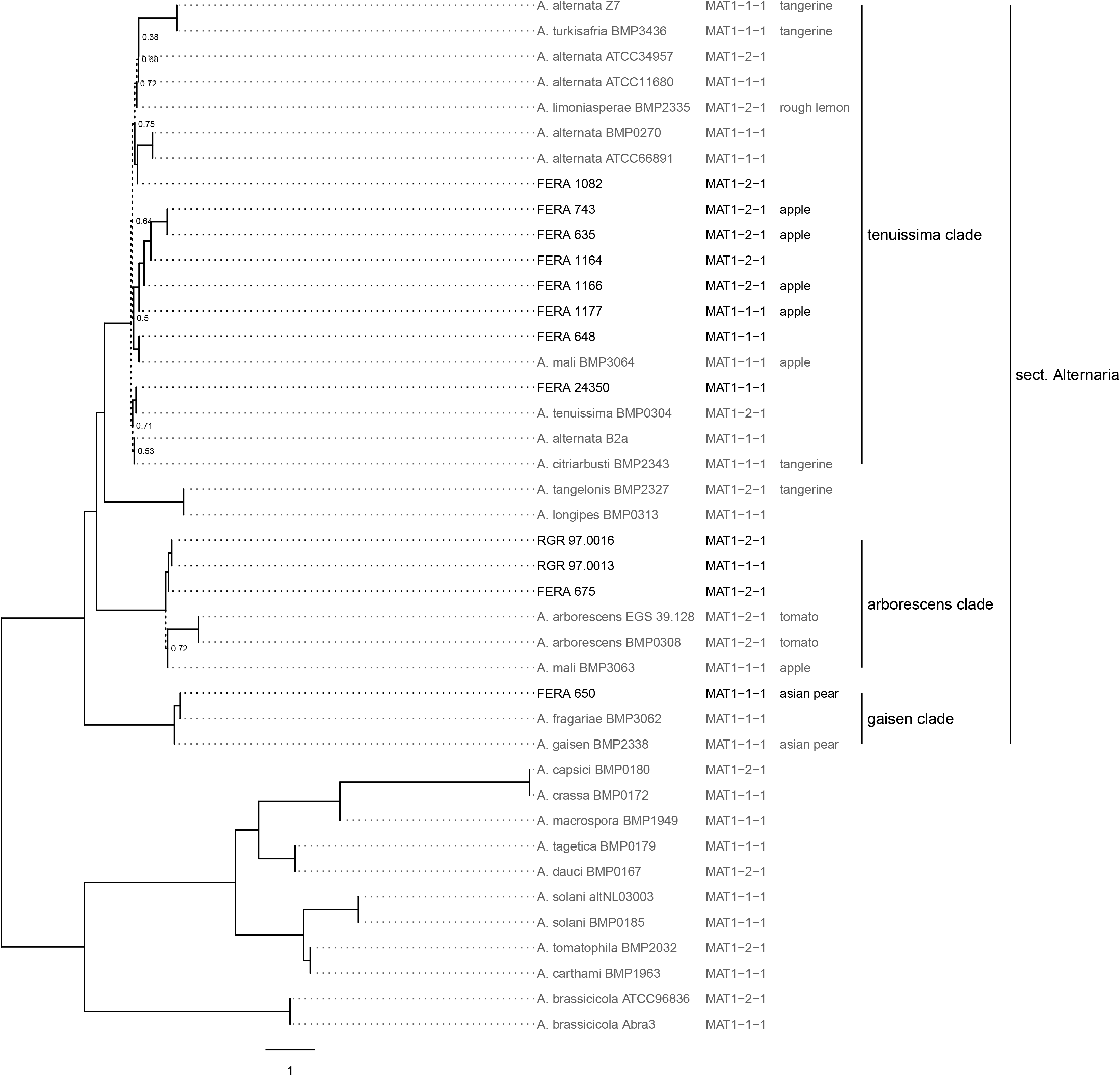
Phylogeny of sequenced and publicly available *Alternaria* spp. genomes. Maximum parsimony consensus phylogeny of 500 conserved single copy loci. Dotted lines show branches with support from < 80% of trees. Mating type idiomorphs MAT1-1-1 and MAT1-2-1 show distribution throughout the phylogeny. Isolates pathotype is labelled following identification of genes involved in synthesis of apple, pear, strawberry, tangerine, rough lemon and tomato toxins.

### 3.3 Gene and effector identification

Gene prediction resulted in 12757 – 13733 genes from the our assemblies (Table 2), with significantly more genes observed in the apple pathotype isolates than in *A. tenuissima* clade non-pathotype isolates (P > 0.01, F_2,8df_ = 51.19). BUSCO analysis identified that gene models included over 97% of the single copy conserved ascomycete genes, indicating well trained gene models. Apple pathotype isolates possessed greater numbers of secondary metabolite clusters (P > 0.01, F_2,8df_ = 8.96) and secreted genes (P > 0.01, F_2,8df_ = 44.21) than non-pathotype *A. tenuissima* isolates, indicating that CDCs contain additional secreted effectors. Non-pathotype *A. tenuissima* clade isolates were found to possess greater numbers of genes encoding secreted proteins than *A. arborescens* isolates (P > 0.01, F_2,8df_ = 44.21), including secreted CAZYmes (P > 0.01, F_2,8df_ = 9.83). The basis of differentiation between these taxa was investigated further.

### 3.4 Genomic differences between *A. tenuissima* and *A. arborescens* clades

Orthology analysis was performed upon the combined set of 158,280 total proteins from the 12 sequenced isolates. In total, 99.2 % of proteins clustered into 14,187 orthogroups. Of these, 10,669 orthogroups were shared between all isolates, with 10,016 consisting of a single gene from each isolate. This analysis allowed the identification of 239 orthogroups that were either unique to *A. arborescens* isolates or expanded in comparison to non-pathotype *A. tenuissima* isolates.

Expanded and unique genes to *A. arborescens* isolates was further investigated using *FERA 675* (Supp. information 2). Genes involved in reproductive isolation were in this set, including 21 of the 148 heterokayon incompatibility (HET) loci from *FERA 675*. CAZymes were also identified within this set, three of which showed presence of chitin binding activity and the other three having roles in xylan or pectin degradation. In total, 25 genes encoding secreted proteins were within this set, secreted proteins with pathogenicity-associated functional annotations included a lipase, a chloroperoxidase, an aerolysin-like toxin, a serine protease and an aspartic peptidase. A further six secreted genes had an effector-like structure by EffectorP but no further functional annotations. Furthermore, one gene from this set was predicted to encode a fungal-specific transcription factor unique to *A. arborescens* isolates.

Further to the identification of genes unique or expanded in *A. arborescens*, 220 orthogroups were identified as unique or expanded in the *A. tenuissima*. These orthogroups were further investigated using isolate *FERA 648* (Supp. information 2). This set also contained genes involved in reproductive isolation, including nine of the 153 from *FERA 648*. CAZymes within the set included two chitin binding proteins, indicating a divergence of LysM effectors between *A. tenuissima* and *A. arborescens* lineages. The five additional CAZymes in this set represented distinct families from those expanded/unique in *A. arborescens*, including carboxylesterases, chitooligosaccharide oxidase and sialidase. In total, 18 proteins from this set were predicted as secreted, including proteins with cupin protein domains, leucine rich-repeats, astacin family peptidase domains and with four predicted to have effector-like structures but no further annotations. *A. tenuissima* isolates had their own complement of transcription factors, represented by four genes within this set.

### 3.5 Identification of CDC contigs and assessment of copy number

Alignment of Illumina reads to the apple and Asian pear pathotype MinION reference assemblies identified variable presence of some contigs, identifying these as contigs representing CDCs (CDC contigs). Six contigs totaling 1.87 Mb were designated as CDCs in the apple pathotype reference (Table 3) and four contigs totaling 1.47 Mb designated as CDCs in the pear pathotype reference (Table 4). Two *A. tenuissima* clade non-pathotype isolates (*FERA 1082*, *FERA 1164*) were noted to possess apple pathotype CDC contigs 14 and 19 totaling 0.78 Mb (Table 3).

**Table 3.** Identification of CDC regions in the *A. alternata* apple pathotype reference genome. Read depth is shown from illumina reads against each contig, by isolate. Contigs showing reductions in coverage from non-pathotype isolates were identified as regions of conditionally dispensable chromosomes (CDCs).

| FERA<br>1166<br>contig | Length<br>(bp) | *A. gaisen* | *A. tenuissima* |  |  |  |  |  |  |  | *A. arborescens* |  |  |
| --- | --- | --- | --- | --- | --- | --- | --- | --- | --- | --- | --- | --- | --- |
|  |  | pear | apple |  |  |  |  |  |  |  | - | - | - |
|  |  | FERA 650 | FERA 1166 | FERA 635 | FERA 743 | FERA 1177 | FERA 648 | FERA 1082 | FERA 1164 | FERA 24350 | FERA 675 | RGR 97.0013 | RGR 97.0016 |
| 1 | 3902980 | 32 | 49 | 26 | 40 | 60 | 34 | 21 | 28 | 36 | 53 | 35 | 26 |
| 2 | 3781932 | 31 | 49 | 26 | 40 | 58 | 33 | 20 | 29 | 35 | 51 | 34 | 26 |
| 3 | 3000390 | 33 | 50 | 26 | 40 | 59 | 34 | 21 | 28 | 36 | 53 | 35 | 26 |
| 4 | 2851745 | 32 | 49 | 27 | 41 | 59 | 34 | 21 | 28 | 36 | 53 | 36 | 27 |
| 5 | 2693844 | 33 | 49 | 27 | 40 | 60 | 34 | 21 | 29 | 36 | 53 | 36 | 27 |
| 6 | 2583941 | 33 | 50 | 27 | 40 | 60 | 34 | 21 | 29 | 36 | 53 | 36 | 27 |
| 7 | 2502671 | 33 | 49 | 26 | 40 | 59 | 34 | 21 | 28 | 36 | 52 | 35 | 26 |
| 8 | 2455819 | 32 | 49 | 27 | 40 | 60 | 33 | 21 | 29 | 35 | 52 | 35 | 26 |
| 9 | 2451092 | 32 | 49 | 26 | 40 | 59 | 33 | 20 | 28 | 35 | 52 | 35 | 26 |
| 10 | 2402550 | 31 | 49 | 26 | 40 | 58 | 32 | 21 | 28 | 35 | 51 | 34 | 25 |
| 11 | 2194253 | 32 | 49 | 26 | 40 | 59 | 34 | 21 | 28 | 35 | 52 | 35 | 26 |
| 12 | 1303685 | 33 | 49 | 27 | 40 | 59 | 34 | 21 | 28 | 36 | 53 | 36 | 27 |
| 13 | 767820 | 23 | 48 | 24 | 36 | 47 | 25 | 17 | 25 | 29 | 44 | 26 | 20 |
| 14 (CDC) | 549494 | 0 | 81 | 29 | 79 | 115 | 0 | 23 | 33 | 0 | 0 | 0 | 0 |
| 15 (CDC) | 435297 | 62 | 90 | 51 | 79 | 407 | 0 | 0 | 0 | 0 | 0 | 0 | 0 |
| 16 | 433285 | 23 | 48 | 26 | 40 | 53 | 26 | 18 | 28 | 33 | 51 | 27 | 20 |
| 17 | 391795 | 26 | 48 | 20 | 33 | 56 | 30 | 19 | 27 | 29 | 51 | 30 | 23 |
| 18 (CDC) | 368761 | 0 | 49 | 22 | 40 | 32 | 0 | 0 | 0 | 0 | 0 | 0 | 0 |
| 19 (CDC) | 225742 | 0 | 48 | 22 | 61 | 72 | 0 | 18 | 31 | 0 | 0 | 0 | 0 |
| 20 (CDC) | 154051 | 0 | 69 | 20 | 45 | 214 | 0 | 0 | 0 | 0 | 0 | 0 | 0 |
| 21 (CDC) | 143777 | 0 | 49 | 20 | 36 | 219 | 0 | 0 | 0 | 0 | 0 | 0 | 0 |
| 22 | 109256 | 25 | 50 | 20 | 30 | 50 | 26 | 14 | 22 | 30 | 35 | 27 | 22 |

**Table 4.** Identification of CDC regions in the *A. alternata* Asian pear pathotype reference genome. Read depth is shown from illumina reads against each contig, by isolate Contigs showing reductions in coverage from non-pathotype isolates were identified as regions of conditionally dispensable chromosomes (CDCs).

| FERA<br>650<br>contig | Length<br>(bp) | *A. gaisen*<br>pear | *A. tenuissima*<br>apple |  |  |  |  |  |  |  | *A. arborescens*<br>- |  |  |
| --- | --- | --- | --- | --- | --- | --- | --- | --- | --- | --- | --- | --- | --- |
|  |  | FERA 650 | FERA 1166 | FERA 635 | FERA 743 | FERA 1177 | FERA 648 | FERA 1082 | FERA 1164 | FERA 24350 | FERA 675 | RGR 97.0013 | RGR 97.0016 |
| 1 | 6257968 | 35 | 43 | 25 | 38 | 58 | 33 | 20 | 28 | 35 | 50 | 34 | 26 |
| 2 | 2925786 | 35 | 45 | 25 | 39 | 58 | 34 | 20 | 28 | 35 | 51 | 35 | 26 |
| 3 | 2776589 | 35 | 44 | 26 | 39 | 58 | 34 | 20 | 28 | 35 | 51 | 35 | 26 |
| 4 | 2321443 | 35 | 45 | 26 | 39 | 58 | 34 | 20 | 28 | 35 | 51 | 35 | 27 |
| 5 | 2116911 | 35 | 46 | 26 | 40 | 59 | 35 | 21 | 29 | 36 | 52 | 36 | 27 |
| 6 | 2110033 | 35 | 44 | 25 | 39 | 58 | 33 | 20 | 28 | 35 | 51 | 35 | 26 |
| 7 | 1975041 | 35 | 42 | 25 | 38 | 56 | 33 | 19 | 27 | 34 | 50 | 34 | 26 |
| 8 | 1822907 | 35 | 43 | 25 | 39 | 58 | 33 | 20 | 28 | 35 | 51 | 35 | 26 |
| 9 | 1811374 | 34 | 42 | 25 | 38 | 57 | 32 | 19 | 27 | 34 | 49 | 34 | 25 |
| 10 | 1696734 | 34 | 42 | 25 | 38 | 56 | 32 | 19 | 27 | 34 | 50 | 34 | 26 |
| 11 | 1617057 | 35 | 43 | 25 | 38 | 57 | 33 | 19 | 27 | 34 | 50 | 34 | 26 |
| 12 | 1416541 | 35 | 47 | 26 | 40 | 60 | 35 | 21 | 29 | 36 | 53 | 36 | 27 |
| 13 | 1110254 | 35 | 39 | 24 | 36 | 55 | 31 | 19 | 26 | 33 | 48 | 32 | 24 |
| 14 (CDC) | 629968 | 68 | 49 | 21 | 41 | 202 | 0 | 0 | 0 | 0 | 0 | 0 | 0 |
| 15 | 554254 | 34 | 40 | 24 | 37 | 56 | 32 | 19 | 27 | 33 | 48 | 33 | 25 |
| 16 (CDC) | 547262 | 34 | 8 | 13 | 18 | 22 | 8 | 5 | 14 | 11 | 20 | 8 | 6 |
| 17 | 522351 | 35 | 37 | 23 | 35 | 55 | 31 | 18 | 26 | 33 | 47 | 33 | 24 |
| 18 | 463030 | 34 | 30 | 22 | 33 | 49 | 25 | 17 | 24 | 29 | 44 | 28 | 21 |
| 19 | 353531 | 34 | 30 | 21 | 34 | 49 | 26 | 16 | 23 | 30 | 43 | 28 | 21 |
| 20 | 350965 | 34 | 20 | 18 | 27 | 45 | 22 | 15 | 19 | 26 | 40 | 20 | 18 |
| 21 | 313768 | 35 | 24 | 19 | 29 | 46 | 25 | 15 | 22 | 27 | 38 | 28 | 19 |
| 22 | 288922 | 33 | 35 | 23 | 35 | 51 | 28 | 21 | 24 | 31 | 44 | 30 | 22 |
| 23 (CDC) | 206183 | 34 | 1 | 3 | 5 | 43 | 0 | 0 | 10 | 1 | 0 | 0 | 0 |
| 24 (CDC) | 89891 | 70 | 0 | 0 | 0 | 0 | 0 | 0 | 0 | 0 | 0 | 0 | 0 |
| 25 | 25201 | 16 | 0 | 0 | 0 | 0 | 0 | 0 | 0 | 0 | 0 | 0 | 0 |
| 26 | 23394 | 5 | 0 | 0 | 0 | 0 | 0 | 0 | 0 | 0 | 0 | 0 | 0 |
| 27 | 19592 | 17 | 0 | 0 | 0 | 0 | 0 | 0 | 0 | 0 | 0 | 0 | 0 |

Read alignments showed that CDC contigs were present in multiple copies within *A. alternata* pathotype isolates. *FERA 1166* Illumina reads aligned to its own assembly showed two-fold coverage over contigs 14, 15, 20 and 21 in comparison to core contigs (Table 3). This was more pronounced in isolate *FERA 1177* that had between two- and eight-fold coverage of these contigs. The same was observed in pear pathotype CDC regions, with contigs 14 and 24 in isolate *FERA 650* showing two-fold coverage from Illumina reads in comparison to core contigs (Table 4).

### 3.6 Toxin gene clusters are present on multiple CDC contigs

Homologs to 15 of the 17 AMT cluster genes were located on contigs 20 and 21 in the apple pathotype reference genome (e-value < 1×10^−30^, > 70% query alignment), confirming them as CDC-regions (Table 5). Of the remaining two genes, *AMT11* had low-confidence BLAST homologs on contigs 18 and 21 (e-value < 1×10^−30^) whereas the best BLAST hit of *AMT15* was located on contig 18 (e-value < 1×10^−30^). Duplication of toxin gene regions was observed between CDC contigs, with contig 20 carrying homologs to 16 toxin genes, but with contig 21 also carrying the *AMT1* to *AMT12* section of the cluster (Table 5). The three other apple pathotype isolates (*FERA 635*, *FERA 743* and *FERA 1177*) also showed presence of 15 of the 17 AMT genes (e-value < 1×10^−30^, > 70% query alignment), and with some AMT genes present in multiple copies within the genome indicating that the AMT toxin region has also been duplicated in these isolates.

**Table 5.**
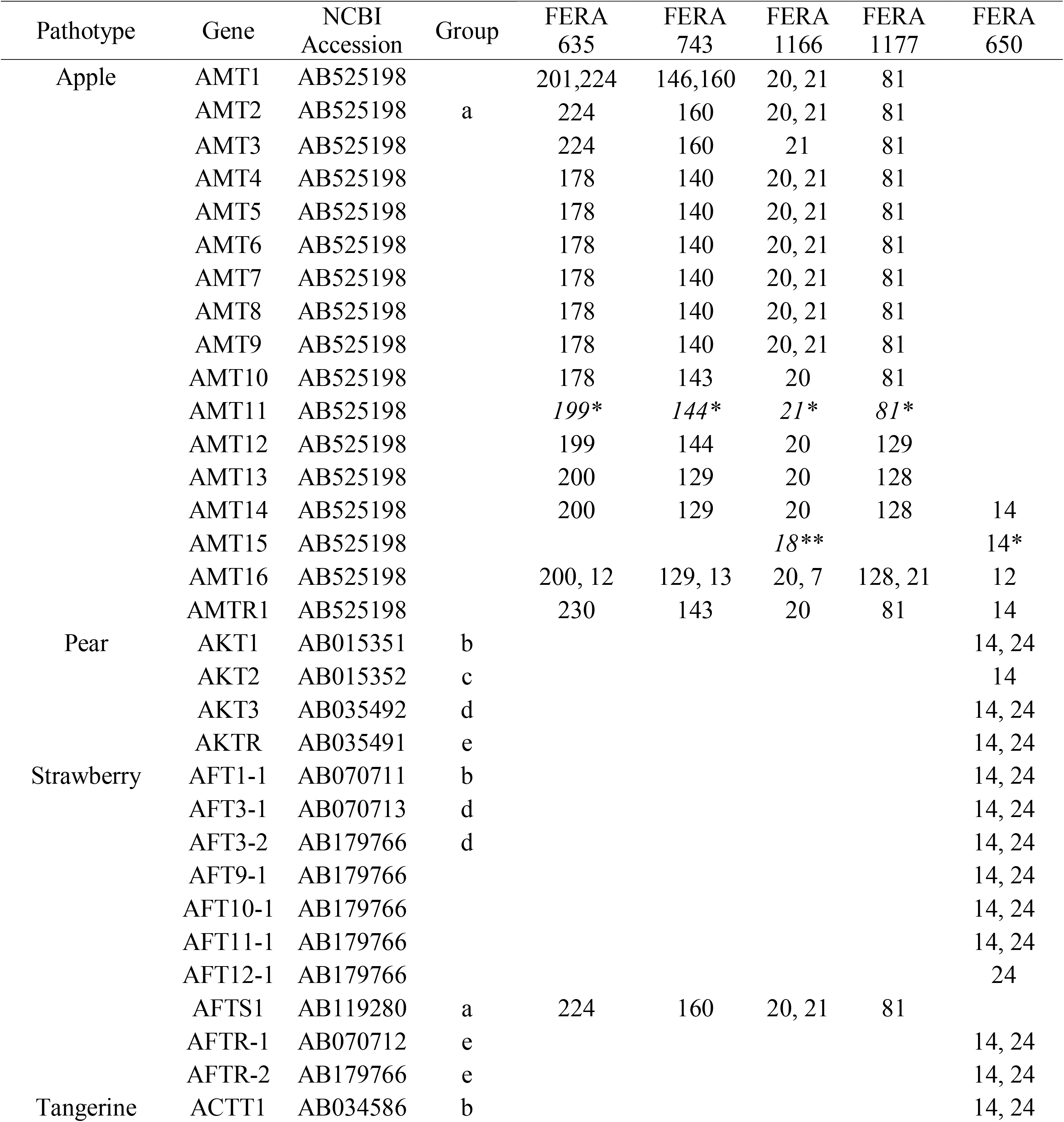
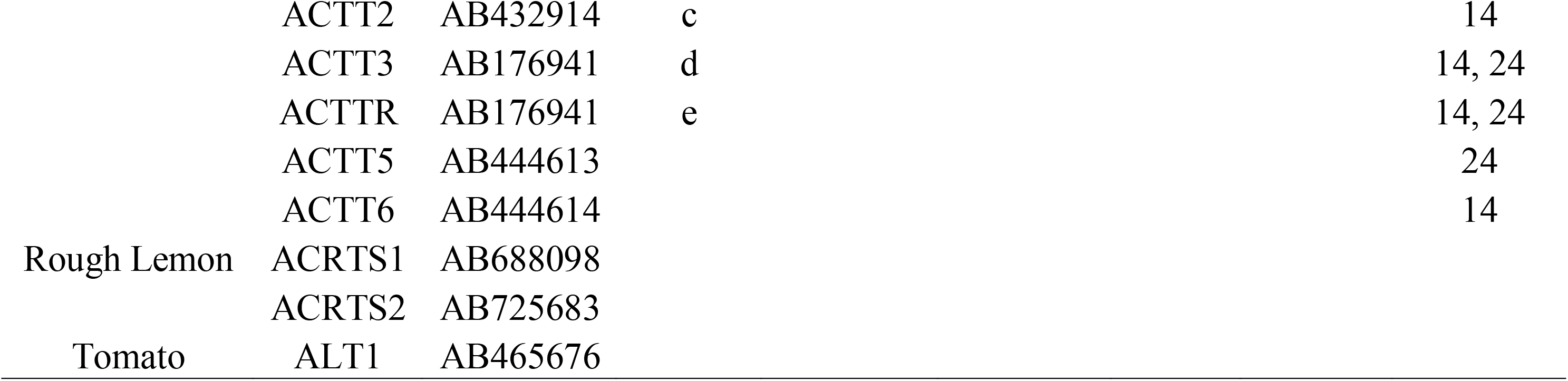
Genomic location (contig number) of homologs to genes from apple, pear, strawberry, tangerine, rough lemon, and tomato toxin gene clusters. Results from reference genome isolates *FERA 1166* and *FERA 650* indicate toxin clusters are present in multiple copies within the genome. This is supported by identification of multiple AMT1 homologs in *FERA 635* and *FERA 743*. Homology between query sequences is shown (homolog groups), as determined from reciprocal BLAST searches between queries. Homologs are identified by e-value < 1×10^−30^, > 70% query alignment. * Marks lower-confidence hits with < 70% query alignment.

The Asian pear pathotype was also found to carry toxin gene clusters in multiple copies, with homologs to the four AKT cluster genes present on contig 14 of the *FERA 650* assembly (e-value < 1×10^−30^, > 70% query alignment), with three of these also present on contig 24 (e-value < 1×10^−30^, two with > 70% query alignment). BLAST hit results from AKT genes were supported by their homologs from strawberry and tangerine pathotypes also found in these regions (Table 5). The pear pathotype genome was also found to contain additional homologs from apple (*AMT14*), strawberry (*AFT9-1*, *AFT10-1*, *AFT11-1* and *AFT12-1*) and citrus (*ACTT5* and *ACTT6*) located on CDC contigs 14 and 24 (Table 5).

### 3.7 CDCs carry effectors alongside secondary metabolites

A total of 624 proteins were encoded on the six contigs designated as CDCs in the reference apple pathotype genome, with 502 proteins encoded on the four Asian pear pathotype CDC contigs (Supp. information 3). We further investigated the gene complements of these regions.

Approximately a quarter of gene models on apple pathotype CDC contigs were involved in secondary metabolism, with 153 genes present in six secondary metabolite gene clusters. This included AMT toxin gene homologs on contigs 20 and 21, which were located within NRPS secondary metabolite gene clusters. Three other secondary metabolite clusters were located on CDC contigs with two of these involved in the production of T1PKS secondary metabolites and the third with unknown function. A further two secondary metabolite clusters were located on contig 14 shared with two non-pathotype isolates, one of which is involved in the production of a T1PKS. The pear pathotype also carried 153 genes in secondary metabolite gene clusters. These 30 % of CDC genes were located in four clusters, with the AKT toxin genes in T1PKS clusters of contigs 14 and 24. A second cluster was present on contig 14 with unknown function and a T1PKS cluster was present on contig 16.

Approximately 5 % of the genes on apple CDC contigs encoded secreted proteins, with 32 in isolate *FERA 116*6 many of which had potential effector functions with six designated as CAZymes and 12 testing positive by EffectorP. Similarly, a total of 41 secreted proteins were predicted on the CDC regions of the Asian pear pathotype, with eight of these designated as secreted CAZymes and 13 testing positive by EffectorP. Further investigation into the 32 secreted proteins from the apple pathotype identified three CAZYmes from the chitin-active AA11 family, two from the cellulose-active GH61 family and one cellulose-active GH3 family protein. Six of the 13 EffectorP proteins also had domains identifiable by interproscan: four carried NTF2-like domains, which are envelope proteins facilitating protein transport into the nucleus; one was a fungal hydrophobin protein; one was a member of an panther superfamily PTHR40845 that shares structural similarity with proteins from the plant pathogens *Phaeospharia nodorum*, *Sclerotinia sclerotiorum* and *Ustilago maydis*. Of the 38 secreted proteins identified from the pear pathotype, two CAZYmes were also identified from the chitin-active AA11 family, two from the AA3 family with single proteins from GH5, CBM67 and AA7 families. Ten of the twelve secreted EffectorP proteins had no functional information as predicted by interproscan, with the other two identified as carrying WSC domains IPR002889, which are cysteine-rich domains involved carbohydrate binding. CDCs may also play important roles in transcriptional regulation with 29 putative transcription factors identified in the apple pathotype CDC contigs(4.6 % CDC genes) and 35 identified in pear pathotype CDC contigs (7.0 % CDC genes).

### 3.8 Polyphyletic distribution of apple and tangerine pathotypes

The evolutionary relationship between *A. alternata* pathotypes sequenced in this study and publicly available genomes was analyzed by the core gene phylogeny (Fig. 2). We identified four isolates as tangerine pathotypes (Z7, BMP2343, BMP2327, BMP3436) two as tomato pathotypes (BMP0308, EGS39-128), one Asian pear pathotype (MBP2338), one rough-lemon (BMP2335) and two apple pathotypes (BMP3063, BMP3064) through searches for genes from HST-gene clusters (Supp. information 4). When plotted on the genome phylogeny, we found the apple and tangerine pathotypes to be polyphyletic (Fig. 2). Five of the six sequenced apple pathotype isolates were located in the *A. tenuissima* clade and one in the *A. arborescens* clade, whereas the tangerine pathotype was present in both the *A. tenuissima* clade and in the *A. tangelonis / A. longipes* clade.

### 3.9 Molecular tools for identification of apple, pear and strawberry pathotypes

PCR primers for three loci *(AMT4*, *AKT3* and *AMT14*) were designed to identify the distribution of pathotypic isolates through the *A. alternata* species group and were screened against a set of 89 previously characterised isolates (Fig. 4). Five isolates tested positive for the presence of *AMT4*, each of which was from the *A. tenuissima* clade (*FERA 635*, *FERA 743*, *FERA 1166*, *FERA 1177*). Five isolates tested positive for the presence of *AKT3*, including the three isolates from Asian pear in the *A. gaisen* clade and a further two isolates from the *A. tenuissima* clade that were from strawberry. Sequencing of the *AKT3* amplicons from the two isolates were strawberry identified them as the *AFT3-2* ortholog of *AKT3*, showing that these isolates were strawberry pathotypes rather than pear pathotypes. Sequencing of PCR products from the other isolates confirmed them to be apple or pear pathotypes as expected. All of the isolates testing positive for *AMT4* or *AKT3* also tested positive for *AMT14*, indicating its suitability as a target gene for identification of a range of pathotypes.

**Figure 4.**
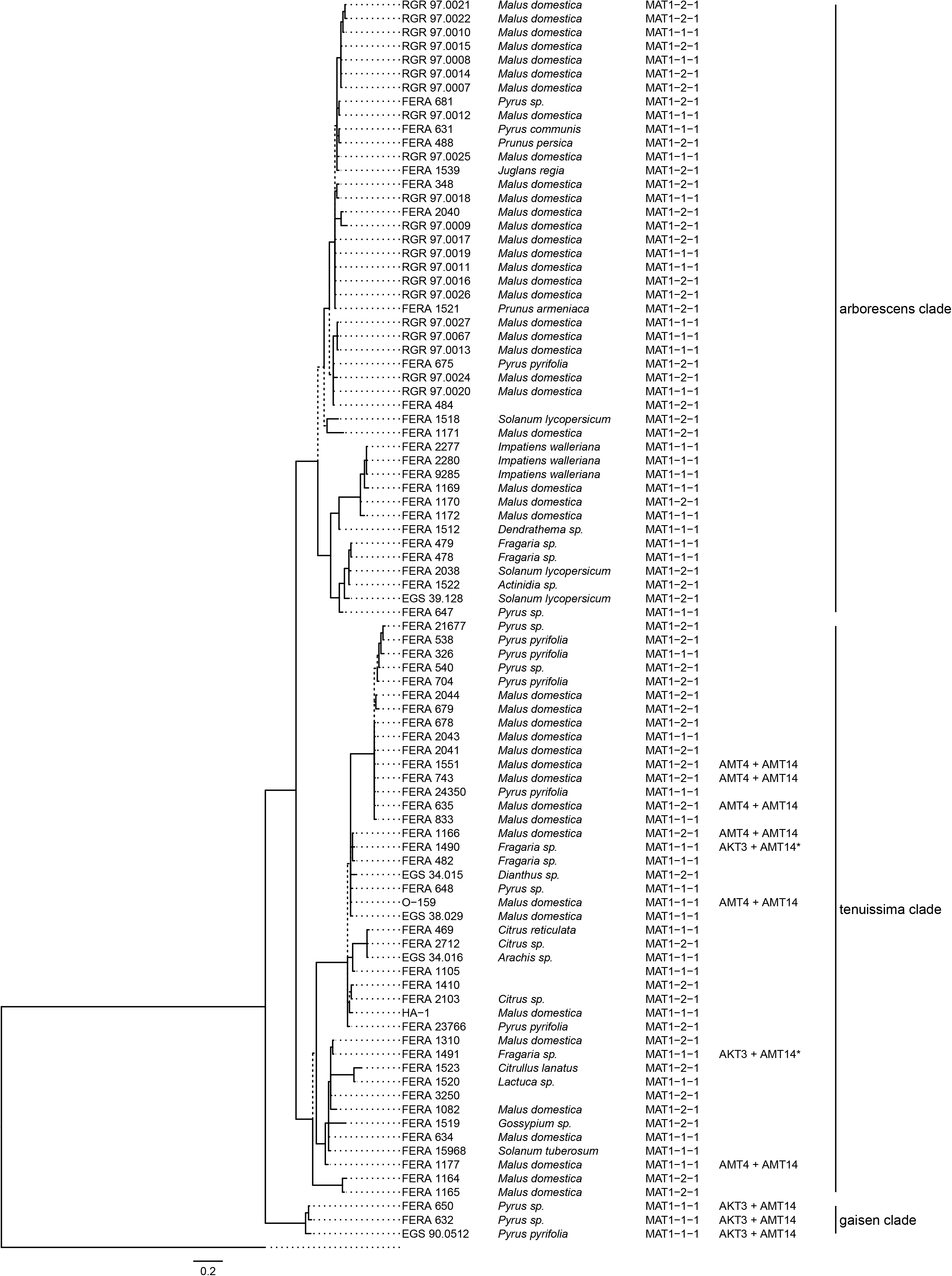
Presence/absence of toxin and mating type genes for 89 *Alternaria* isolates. Results are plotted onto the 5-gene phylogeny of Armitage (2015). MAT1-1-1 and MAT1-2-1 mating type idiomorphs are designated.

Presence of apple pathotype CDCs was confirmed to be associated with pathogenicity through detached apple leaf assays. Apple pathotype isolates showed significantly greater numbers of necrotic lesions when inoculated onto *cv.* Spartan (F_72df_ =100.64) and *cv.* Bramley’s Seedling (F_72df_ =69.64) leaves than non-pathotype *A. tenuissima* isolates (Fig. 5).

**Figure 5.**
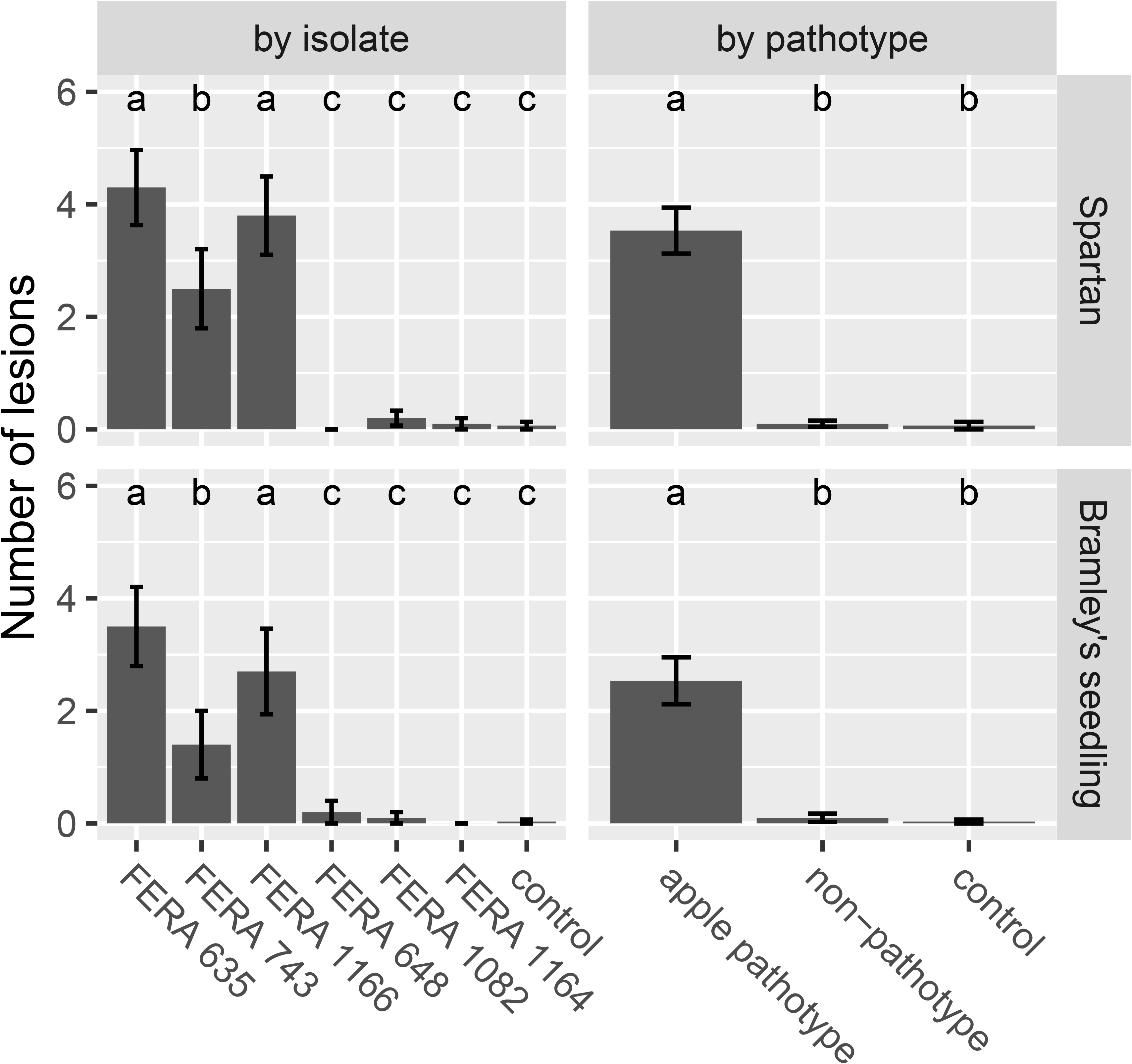
Number of *Alternaria* lesions per apple leaf at 14 dpi for treatments in virulence assays on *cv.* Spartan or *cv.* Bramley’s Seedling leaves. Apple pathotype isolates (*FERA 635*, *FERA 743* and *FERA 1166*) cause significant disease symptoms in comparison to control leaves, in contrast to non-pathotype isolates (*FERA 648*, *FERA 1082* and *FERA 1164*). Number of lesions (±SE) are shown with significance at P < 0.05, as determined from a GLM at the isolate and pathotype level.

### 3.10 Signatures of genetic exchange

Of the 12 sequenced isolates, BLAST searches identified five as carrying the MAT1-1-1 idiomorph and seven as carrying MAT1-2-1. Both idiomorphs showed distribution throughout the *A. alternata* genome phylogeny (Fig. 2). These results were supported by PCR assays identifying the mating type of 89 previously characterised isolates (Fig. 4). Idiomorphs did not deviate from a 1:1 ratio within *A. tenuissima* (21 MAT1-1-1 : 23 MAT1-2-1; χ^2^ = 0.09, 1df, P>0.05) or *A. arborescens* clades (18 MAT1-1-1 : 24 MAT1-2-1; χ^2^ = 0.86; 1df; P>0.05), as expected under a random mating population. All three of the *A. gaisen* clade isolates carried the MAT1-2-1 idiomorph.

## 4 Discussion

This work builds upon the current genomic resources available for *Alternaria*, including the *A. brassicicola* and *A. solani* genomes (Belmas et al. 2018; Wolters et al. 2018), *A. alternata* from onion (Bihon et al. 2016) the additional 25 *Alternaria spp.* genomes available on the *Alternaria* Genomes Database (Dang et al. 2015) as well as recent genomes for other pathotype and non-pathotype *A. alternata* (Hou et al. 2016; Wang et al. 2016; Nguyen et al. 2016). Of the previously sequenced genomes, *A. solani*, the citrus pathotype and a non-pathotype *A. alternata* isolate have benefitted from long read sequencing technology with each comprising less than 30 contigs (Wolters et al. 2018; Wang et al. 2016; Nguyen et al. 2016). Total genome sizes in this study (33-36 Mb) were in line with previous estimates for *A. alternata*, with the tomato pathotype also previously assembled into 34 Mb (Hu et al. 2012). Synteny analysis of our two reference genomes against the chromosome-level *A. solani* genome revealed structural differences for chromosomes 1 and 10 in the apple pathotype and for chromosomes 1 and 2 in the pear pathotype. These structural differences may represent distinct traits between clades of the *A. alternata* species group, and may represent a barrier to genetic exchange involved in the divergence of *A. gaisen* and *A. tenuissima* lineages. The number of essential chromosomes in our reference genomes is in line with previous findings in *A. alternata* (Kodama et al. 1998), with 9-11 core.

Species designations within the species group have been subject to recent revision (Woudenberg et al. 2015; Lawrence et al. 2013; Armitage et al. 2015) leading to potential confusion when selecting isolates for study. For example, the available *A. fragariae* genome (Dang et al. 2015), did not represent a strawberry pathotype isolate and was located in the *A. gaisen* clade. As such, the phylogenetic context for sequenced *Alternaria* genomes described in this study, along with pathotype identification provides a useful framework for isolate selection in future work.

### 4.1 Evidence of genetic exchange

A 1:1 ratio of MAT loci was observed within *A. arborescens* and *A. tenuissima* clades. This supports previous identification of both idiomorphs within *A. alternata*, *A. brassicae* and *A. brassicicola* (Berbee et al. 2003). Furthermore, presence of both MAT idiomorphs within apple pathotype isolates indicates that genetic exchange (sexuality or parasexuality) has occurred since the evolution of CDCs, providing a mechanism of transfer of CDCs. Furthermore, we show that some recent or historic genetic exchange has occurred between *A. tenuissima* and *A. arborescens* clades, with both apple and tangerine pathotypes exhibiting a polyphyletic distribution throughout the phylogeny.

### 4.2 Duplication of toxin-gene contigs

Toxin genes have been proposed to be present in multiple copies within *A.* sect. *alternaria* pathotype genomes with *AMT2* proposed to be present in at least three copies in the apple pathotype CDC (Harimoto et al. 2008), and multiple copies of *AKTR* and *AKT3* in the pear pathotype (Tanaka et al. 1999; Tanaka and Tsuge 2000). Through read mapping we demonstrated that this is the case. Furthermore, we show that toxin gene clusters are present on multiple contigs, with differences in the gene complements between these clusters. At this stage, it is unclear whether these different clusters are responsible for the production of the variant R-groups previously characterised in AMT or AKT toxins (Nakashima et al. 1985; Harimoto et al. 2007). Differences were also noted between non-pathotype isolates from the *A. tenuissima* clade in the presence/absence of contigs 14 and 19, representing a total of 775 Kb. Chromosomal loss has been reported in the apple pathotype (Johnson et al. 2001), and it is not clear if this represents chromosomal instability in culture or additional dispensable chromosomes within *A. tenuissima* clade isolates.

### 4.3 PCR primers for diagnostics

It is now clear that genes on essential chromosomes do not provide reliable targets for identification of different pathotypes and hence loci located directly on CDCs should be used. We found *AMT14* homologs to be present in all pathotype genomes and designed primers to this region. These demonstrated specificity to apple, pear and strawberry pathotypes within a set of 86 *Alternaria* isolates. Furthermore, Sanger sequencing of these amplicons confirmed this to be a single locus that can both identify and discriminate a range of pathotypes. Wider validation of this primer set is now required to test its suitability across other pathotypes.

### 4.4 Divergence of *A. arborescens* and *A. tenuissima*

The divergence of *A. tenuissima* and *A. arborescens* lineages was investigated through identification of expanded and unique gene compliments. We identified HET loci unique to *A. arborescens* or *A. tenuissima* lineages. HET loci may act as incompatibility barriers to common genetic exchange between these taxa (Glass and Kaneko 2003). Taxa also showed divergence in effector profiles, including chitin binding effectors, with *A. arborescens* isolates possessing unique xylan/pectin degradation CAZymes, while *A. tenuissima* isolates possessed unique carboxylesterase, chitooligosaccharide and sialidase CAZymes. Chitin binding proteins are important in preventing MAMP triggered host recognition by plants and animals during infection, and may also aid persistence of resting bodies outside of the host (Kombrink and Thomma 2013). Putative transcription factors were also amongst the proteins specific to *A. arborescens* or *A. tenuissima*, indicating that these taxa not only possess distinct gene complements but also differ in how they respond to stimuli. Dispersed repeat sequences such as transposable elements have been shown to serve as sites of recombination within and between fungal chromosomes (Zolan 1995) and we also show distinct transposon profiles between *A. arborescens* and *A. tenuissima*. Transposons are known to aid host adaptation in plant pathogens (Faino et al. 2016; Gijzen 2009; Schmidt et al. 2013) and have been a mechanism for differentiation of these taxa.

### 4.5 Effectors on CDC regions

*Alternaria* HSTs are capable of inducing necrosis on non-host leaves (Kohmoto et al. 1976), meaning that non-host resistance must be associated with recognition of other avirulence genes. We investigated the complements of other putative pathogenicity genes and effectors produced by the apple and Asian pear pathotypes and identified additional CAZymes and secondary metabolite profiles on CDC regions, distinct between pathotypes, suggesting additional host-adapted tools for pathogenicity. Additional secondary metabolites clusters were present on both apple and pear pathotype CDCs as well as unique complements of secreted CAZymes. CAZyme families AA3, AA7 and AA9 have previously been reported to be in greater numbers in the citrus pathotype in comparison to non-pathotypes (Wang et al. 2016). Furthermore, putative transcription factor genes were identified in CDCs indicating that these regions may have some level of transcriptional autonomy from the core genome. This has been shown in *Fusarium*, where effector proteins are regulated by the SGE transcription factor on the core genome but also by FTF and other transcription factor families (TF1-9) located on lineage specific chromosomes (van der Does et al. 2016).

### 4.6 Conclusions

We report near-complete reference genomes for the apple and Asian pear pathotypes of *A.* sect. *alternaria* and provide genomic resources for a further ten diverse isolates from this clade. For the first time we show sequenced *Alternaria* genomes in a phylogenetic context allowing the identification of both mating type idiomorphs present in *A. arborescens* and *A. tenuissima*, with a distribution throughout subclades that was indicative of recent genetic exchange. The presence of the apple CDC in isolates of both mating types supports gene flow between isolates. Furthermore, the distribution of isolates from different pathotypes throughout the phylogeny indicated that apple and tangerine pathotypes are polyphyletic. This means that gene flow is not limited to within, but has also occurred between *A. tenuissima* and *A. arborescens* lineages. We also developed PCR primers to aid identification of pathotypes, with those targeting the AMT14 locus identifying a range of pathotypes due to its conservation between CDCs. Despite evidence of genetic exchange between *A. arborescens* and *A. tenuissima* clades, we show that these taxa are sufficiently isolated to have diverged, with significant differences in core effector profiles and transposon content.

## Supporting information

Identification of single copy Ascomycete genes in reference Alternaria spp. genomes

Gene IDs, location and functional annotations of genes in expanded orthogroups between A. tenuissima and A. arborescens.

Gene IDs, location and functional annotations of genes located on CDC contigs from apple and Asian pear pathotype isolates.

Toxin gene BLAST hits in publicly available Alternaria spp. genomes, allowing identification of pathotype isolates.

**Supp. information 1**

Identification of single copy Ascomycete genes in reference *Alternaria* spp. genomes

**Supp. information 2**

Gene IDs, location and functional annotations of genes in expanded orthogroups between *A. tenuissima* and *A. arborescens*.

**Supp. information 3**

**Supp. information 4**

Toxin gene BLAST hits in publicly available *Alternaria* spp. genomes, allowing identification of pathotype isolates.

## 6 Data Availability

Accession numbers for genomic data are provided in Table 1. Sanger sequence data is deposited on NCBI under accession numbers MK255031-MK255052.

## 8 Author Contributions

AA, SS, CL and JC contributed conception and design of the study; AA, HC and RH performed lab work including library preparation and sequencing; AA performed bioinformatic analyses and wrote the manuscript. All authors contributed to manuscript revision, read and approved the submitted version.

## 10 Acknowledgments

Thanks are given to the FERA Science Ltd, Dr. P. Gannibal, Dr. R. Roberts and Dr. E. Simmons for access to *Alternaria* isolates. This manuscript has been released as a Pre-Print on bioRxiv (Armitage et al. Preprint).

